# Herpes simplex virus entry by a non-conventional endocytic pathway

**DOI:** 10.1101/2020.09.28.317867

**Authors:** Giulia Tebaldi, Suzanne M. Pritchard, Anthony V. Nicola

## Abstract

Herpes simplex virus 1 (HSV-1) causes significant morbidity and mortality in humans worldwide. HSV-1 enters epithelial cells via an endocytosis mechanism that is low pH-dependent. However, the precise intracellular pathway has not been identified, including the compartment where fusion occurs. In this study, we utilized a combination of molecular and pharmacological approaches to better characterize HSV entry by endocytosis. HSV-1 entry was unaltered in both cells treated with siRNA to Rab5 or Rab7 and cells expressing dominant-negative forms of these GTPases, suggesting entry is independent of the conventional endo-lysosomal network. The fungal metabolite brefeldin A (BFA) and the quinoline compound Golgicide A (GCA) inhibited HSV-1 entry via beta-galactosidase reporter assay and impaired incoming virus transport to the nuclear periphery, suggesting a role for trans Golgi network (TGN) functions and retrograde transport in HSV entry. Silencing of Rab9 or Rab11 GTPases, which are involved in the retrograde transport pathway, resulted in only a slight reduction in HSV infection. Together these results suggest that HSV enters host cells by an intracellular route independent of the lysosome-terminal endocytic pathway.

**IMPORTANCE:** HSV-1, the prototype alphaherpesvirus, is ubiquitous in the human population and causes lifelong infection that can be fatal in neonatal and immunocompromised individuals. HSV enters many cell types by endocytosis, including epithelial cells, the site of primary infection in the host. The intracellular itinerary for HSV entry remains unclear. We probed the potential involvement of several Rab GTPases in HSV-1 entry, and suggest that endocytic entry of HSV-1 is independent of the canonical lysosome-terminal pathway. A non-traditional endocytic route may be employed, such as one that intersects with the TGN. These results may lead to novel targets for intervention.

## INTRODUCTION

Herpes simplex viruses (HSVs) are ubiquitous human pathogens that cause lifelong latent infections and significant morbidity and mortality all around the world. In immunocompetent patients HSV can cause cold sores and genital infections. Serious outcomes include blindness and fatal neonatal infections. In immune-compromised persons, HSV infects diverse organ systems including the respiratory and gastrointestinal tracts and the central nervous system (1-3). HSV-1 is internalized by endocytosis into epithelial cells, the site of lytic replication, by a multistep process that requires low pH. HSV enters some cells, including human neurons, by a pH-independent penetration at the plasma membrane (4-9). HSV entry is quite rapid. Immediately after infection, enveloped viral particles are detected in smooth-walled vesicles adjacent to the host cell plasma membrane (10). Treatment of cells with either hypertonic (0.3 M sucrose) medium or with medium that inhibits host cell ATP synthesis specifically blocks both receptor-mediated endocytosis and HSV entry into CHO-receptor and epithelial cells (4). Importantly, these treatments do not interfere with non-endocytic entry into cells that support fusion of herpesvirions with the cell surface (4, 11, 12). The enveloped virion traffics through the host cell vesicular system until it arrives in a compartment of the appropriate low pH, triggering membrane fusion and penetration of the capsid into the cytosol (9, 13, 14). Macropinocytosis-like and phagocytosis-like processes have been implicated (4-6, 10, 15-18). However, the precise intra-vesicular route taken and its regulation remains unclear.

Most animal viruses enter cells by endocytosis, and endosomal low pH is the most common cellular trigger of enveloped virus fusion (19). Inhibitors of vesicle acidification block HSV entry at an early, post-binding step (4, 10). Endosomes are the first acidic compartments in the endocytic pathway (20). Vesicular pH gradually decreases from 6.2 to approximately 5.0 as cargo moves from the early endosome (EE) to the late endosome (LE) and finally to the lysosome (9, 10, 21). The endo-lysosomal system represents a complex and highly dynamic network of interacting and interconnected compartments, which are critical for host cell homeostasis (22). This lysosome-terminal endocytic pathway is a very common route for viruses entering cells via endocytosis (23-26). When cells either lack a required cellular gD-binding receptor for entry or the virus itself is entry-defective, HSV-1 is degraded, presumably in lysosomes (4, 10, 27). We theorized that HSV transits the common lysosome-terminal endocytic pathway during viral entry.

Endocytic trafficking is finely regulated by a large family of small Rab GTPase enzymes. Rab (Ras-related protein in brain) (28) proteins are master regulators of intracellular vesicle transport events, including vesicle formation, actin- and tubulin-dependent vesicle movement, and the interconnection of endosomal and autophagy pathways (29-32). Many pathogens hijack Rab GTPase functions upon invasion (28, 33-36). Rab proteins are involved in several stages of the viral replication cycle, including entry via endocytosis, viral assembly and egress, and viral glycoprotein trafficking (37). Rab5 is located at the cytoplasmic surface of the plasma membrane and on early endosomes. Thus, it is involved in the formation of clathrin-coated vesicles (CCVs), selectively regulating the transport of newly endocytosed vesicles from the plasma membrane to early endosomes. Rab5 mediates the homotypic fusion between early endosomes (38-40). Rab7 is a late endosome-/lysosome-associated small GTPase, the only lysosomal Rab protein identified to date. It regulates the transport from EE to LE and LE to lysosomes. Rab7 also plays an important role in autophagy (41-45). Rab9 is located on the late endosome, and it mediates late endosome-to-trans-Golgi network transport (46, 47). Rab11 is associated with the trans-Golgi network (TGN), post-Golgi vesicles, and recycling compartments. It regulates transport along the recycling pathway, from recycling endosomes to the cell surface, and the retrograde transport from the perinuclear endocytic recycling compartment (ERC) to the TGN (41, 48-51).

Here, a combination of molecular and pharmacological approaches was used to better characterize the mechanism of HSV-1 entry, including the intracellular pathway of incoming HSV. Specifically, we used siRNAs and dominant-negative mutants to investigate the involvement of Rab GTPases in HSV-1 endocytosis. Neither Rab5 nor Rab7 played a major role in HSV-1 endosomal trafficking. Rab9 and Rab11 may play a slight role. Inhibitor results suggest that retrograde transport and TGN function may play important roles during HSV entry by endocytosis.

## RESULTS AND DISCUSSION

### Knockdown of Rab5 or Rab7 does not inhibit HSV infectivity

HSV enters cells by an endocytic pathway that is incompletely characterized. Rab GTPases are central to the conventional lysosome-terminal endocytosis pathway that is utilized by a multitude of cargoes, including entering viruses. Rab5 selectively regulates the transport of newly endocytosed cargo from the plasma membrane to the EE. Rab7 is critical for EE to LE traffic (39, 42). Chinese hamster ovary (CHO) cells expressing a gD-receptor such as herpesvirus entry mediator (HVEM), are well-characterized model cell types for HSV entry by endocytosis (10, 52, 53). CHO-HVEM cells were transfected with siRNA sequences targeting Rab5 or Rab7, or with a scrambled siRNA sequence. Downregulation was verified by Western blot analysis (Fig. 1A). As determined by densitometry, host Rab5 and Rab7 were reduced by 60 and 93%, respectively. Rab downregulation was further confirmed by microscopic analysis of fluorescent transferrin or LysoTracker in siRNA transfected cells. Knockdown of Rab5 or Rab7 altered the subcellular distribution of transferrin and LysoTracker in CHO-HVEM cells (Fig. 2A-B).

**Figure 1.**
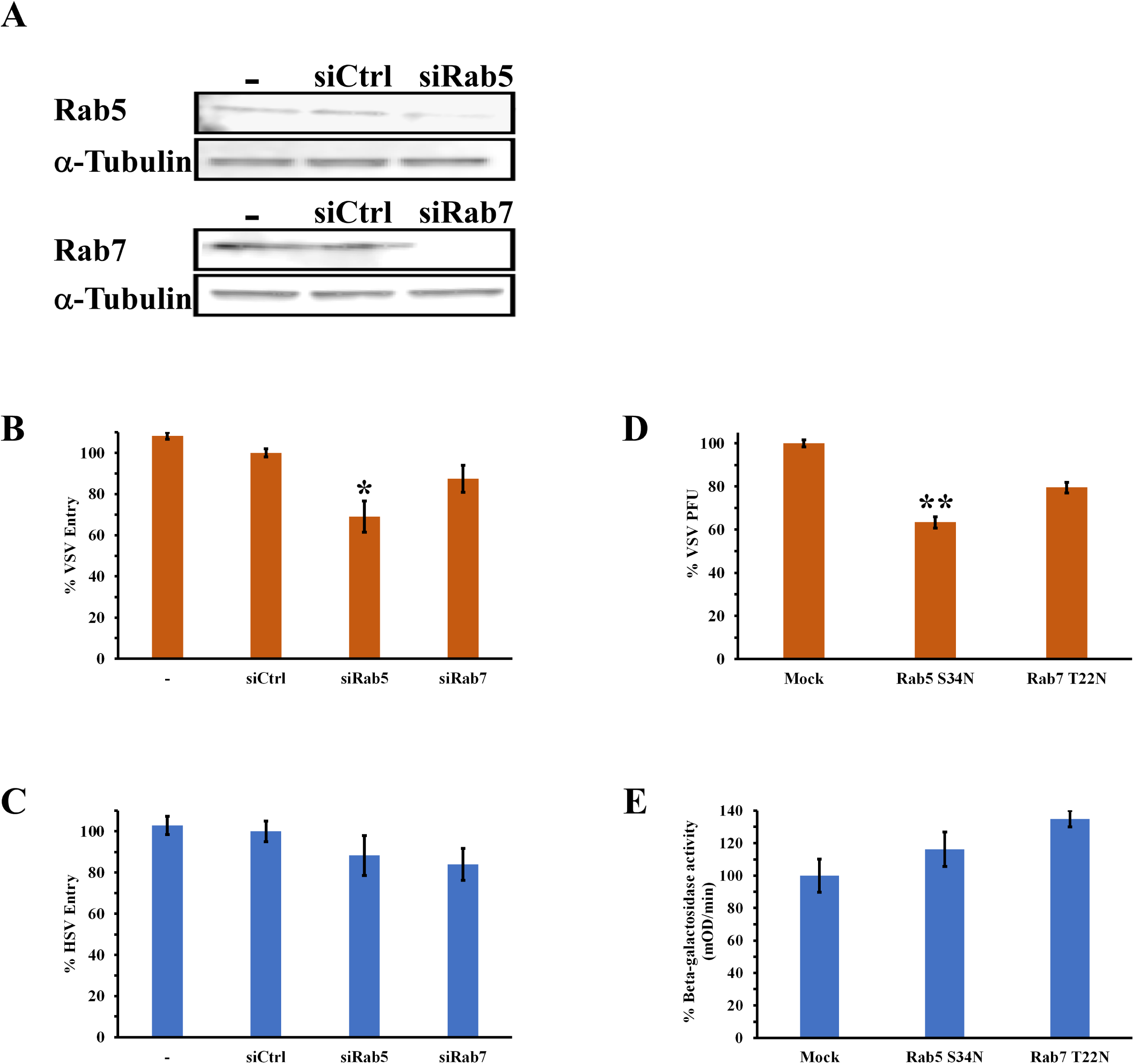
Role of Rab5 and Rab7 on HSV entry and infection. (A) The knockdown efficiency of Rab5 or Rab7 was determined by Western blotting. CHO-HVEM cells (-) were transiently transfected with either control, Rab5 or Rab7 siRNAs for 24 h or 48 h. Cell lysates were reacted with anti-Rab5 or anti-Rab7 antibody specific for the indicated proteins. α-Tubulin was used as an internal loading control. (B, C) CHO-HVEM cells (-) were transiently transfected with control, Rab5 or Rab7 siRNAs for 24 h or 48 h. Control, Rab5, or Rab7 siRNA-treated cells were infected with (B) VSV (MOI of 0.5) or (C) HSV (MOI of 3). At 6 h p.i, infection was detected by quantitating VSV G-positive cells or HSV ICP4-positive cells via indirect immunofluorescence microscopy. At least 500 cells per cover slip were counted. Values are the mean ± SE of three independent experiments *P≤0.05. (D, E) Effect of DN forms of Rab5 or Rab7 on HSV entry. CHO-HVEM cells were stably transfected with control, Rab5 S34N or Rab7 T22N plasmids. Cells were infected with (D) VSV Indiana (120 PFU/well) or (E) HSV-1 KOS (MOI 0.5-1). At 44 h p.i., VSV infectivity was measured by plaque assay. At 6 h p.i., HSV entry was measured by beta-galactosidase reporter assay. Infectivity in control cells was set to 100%. Each experiment was performed in at least triplicate. Data shown are the averages ± SE of at least two independent experiments **P≤0.00005.

**Figure 2.**
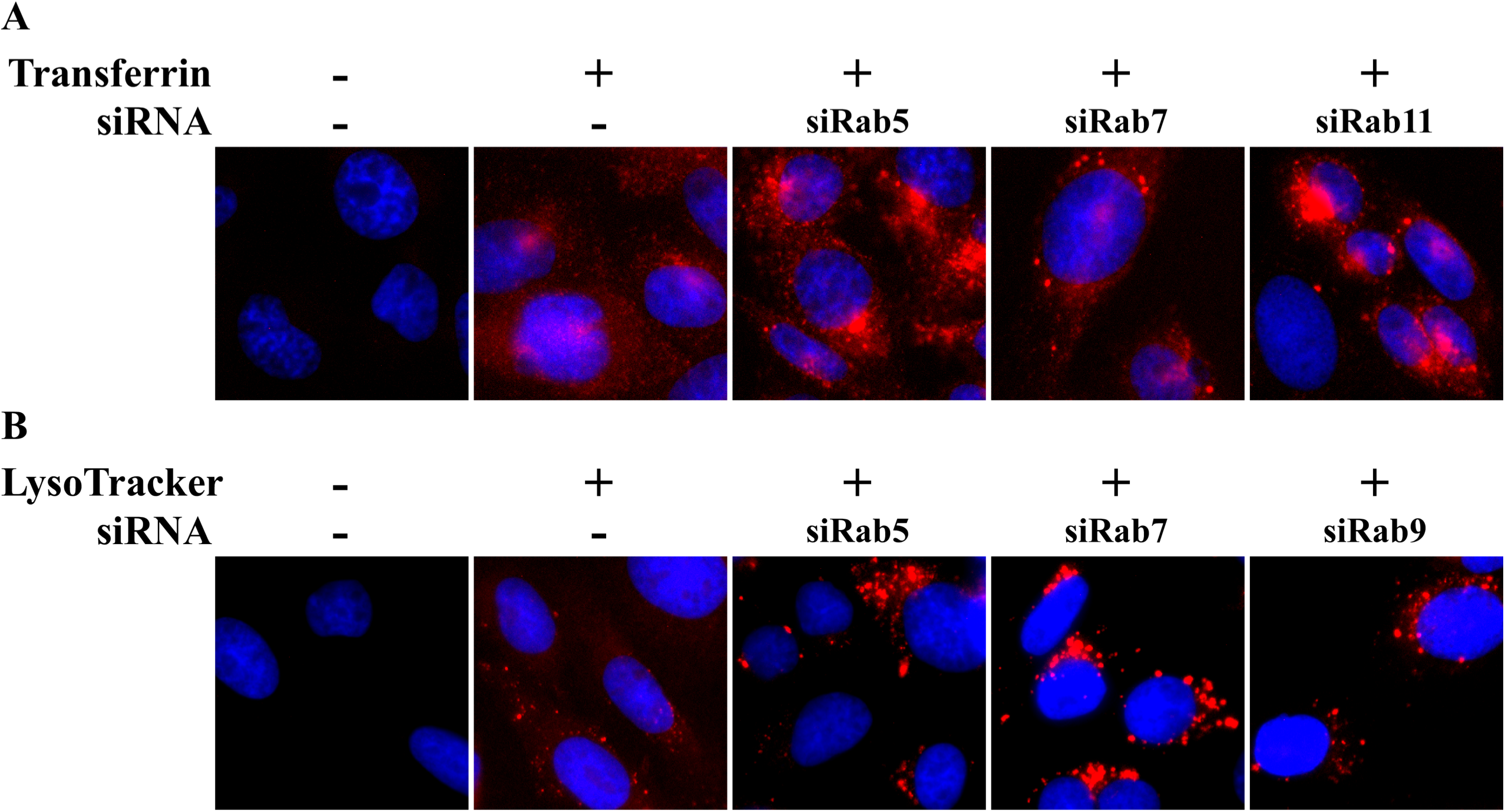
Effect of Rab knockdowns on the subcellular distribution of transferrin and LysoTracker. CHO-HVEM cells were transfected with siRNAs specific for (A, B) Rab5, Rab7, (A) Rab11 or (B) Rab9. Downregulation of Rab5 and Rab7 was verified by Western blot (see Fig. 1). Downregulation of Rab9 and Rab11 was verified by Western blot (see Fig. 4). (A) Transferrin-Texas Red (100 μg/ml) was bound to cells at 4°C, and then cultures were shifted to 37°C for 10 min. (B) 100 nM LysoTracker Red DND-99 was added for 30 min 37°C. Cells were fixed, and nuclei were counterstained with DAPI. The contrast of panels in B was adjusted equivalently with Canvas.

First, the effect of siRNAs was tested on the entry of a control virus, vesicular stomatitis virus (VSV). Endocytic entry of VSV is Rab5-dependent and has also been reported as Rab7-independent (37, 54). The siRNA-transfected cells were infected with VSV Indiana strain for 6 hr (MOI of 0.5). Infectivity was quantitated by indirect immunofluorescence microscopy. As expected, VSV infection of CHO-HVEM cells was inhibited by siRNA to Rab5 (Fig. 1B). Conversely, VSV infection was not significantly inhibited by siRNA to Rab7 (Fig. 1B). These results validate the transfection protocol for down-regulating the production of target proteins in CHO-HVEM cells. Down-regulation of Rab5 resulted in inhibition of infection by a virus that traverses a Rab-5-dependent entry pathway. To address whether HSV entry requires the conventional, lysosome-terminal endosomal pathway, Rab5 or Rab7 was knocked down in CHO-HVEM cells and HSV KOS infectivity at 6 hr p.i. was quantitated. Neither Rab5 nor Rab7 silencing resulted in a significant decrease in HSV-1 infection (Fig. 1C).

To assess further the roles of Rab5 and Rab7 in HSV entry and infection, CHO-HVEM cells were stably transfected with dominant-negative (DN) Rab5 S34N or Rab7 T22N. As expected, VSV infection of Rab5 S34N-expressing cells was reduced relative to control cells, while infection of Rab7 T22N-expressing cells was less affected (Fig. 1D). HSV-1 entry into CHO-HVEM cells, as measured by the reporter assay, was not inhibited by the expression of DN forms of Rab5 or Rab7 (Fig. 1E). Together, the results in Figure 1 suggest that Rab5 and Rab7 GTPases do not play a major role in HSV entry and that virus-cell fusion may occur in an intracellular compartment that is distinct from the conventional endosomal route. The majority of viruses that enter cells via endocytosis use the lysosome-terminal endocytic pathway; however, several viruses have evolved different and sophisticated mechanisms to hijack other aspects of the host endocytic network (23). Lymphocytic choriomeningitis (LCMV) enters cells via LE bypassing the EE (55). Mouse polyomavirus (PyV) travels from the EE to the recycling endosome and then to the ER (56). Papillomavirus takes a retrograde transport pathway to the TGN (57-59).

### Inhibitors of TGN function impair HSV entry

To probe further the involvement of mildly acidic compartments in HSV entry, in the context of a non-conventional vesicular pathway, we utilized pharmacologic inhibitors of the trans-Golgi network, which has a pH of ∽ 6 (60, 61). The pH of the CHO cell TGN is 5.95 (62), which is consistent with the pH that induces conformational change in HSV-1 gB (13, 63-66). Brefeldin A (BFA) is a small hydrophobic compound produced by toxic fungi (67). It triggers the absorption of the cis/medial/trans portion of the Golgi apparatus (68) into the ER through inhibition of the cis-Golgi ArfGEF (guanine nucleotide exchange factor) (GBF1). GBF1 is responsible for maintaining Golgi structure and enabling anterograde and retrograde traffic through the Golgi and TGN (69). BFA’s effects are not limited to the Golgi. BFA also promotes the tubulation of early endosomes, the lysosome and TGN (67) whose components redistribute with the recycling endosomal system (67, 70, 71). Golgicide A (GCA) is a potent and highly specific inhibitor of GBF1. Inhibition of GBF1 function arrests the retrograde transport of Shiga toxin from EE/RE to the TGN (69, 72).

The effect of BFA and GCA on HSV-1 entry was measured by a beta-galactosidase reporter assay. CHO-HVEM cells harbor the *lacZ* gene under an HSV-inducible promoter. BFA and GCA inhibited HSV-1 entry into CHO-HVEM cells in a concentration-dependent manner by as much as 46% and 79%, respectively (Fig. 3A, B). To extend these results, we investigated the role of the TGN in delivery of incoming HSV-1 K26GFP capsids to the nucleus during entry (Fig. 3C). In HSV-infected CHO-HVEM cells, the bulk of the GFP signal is detected at or near the nucleus by 2.5 h p.i (Fig. 3C). In the continued presence of BFA or GCA, GFP-tagged capsids were not effectively transported to the nuclear periphery (Fig. 3C). Instead, the bulk of GFP-tagged viral particles appeared to be trapped at sites distinct from the nucleus. These results were obtained in the presence of the protein synthesis inhibitor cycloheximide to ensure that GFP signal was from input virions only. This also suggests that newly synthesized viral factors do not affect the TGN-dependence of entry. Together, the results suggest that the TGN and the retrograde transport pathway play an important role during HSV entry by endocytosis. Nonenveloped viruses such as simian virus 40 (SV40), human polyomaviruses (73), adeno-associated virus (74, 75), and human papillomavirus (HPV) (57-59), take advantage of the retrograde transport route during entry. HPV entry is inhibited by BFA at a post-fusion step (76). The possibility remains that BFA inhibits a post-fusion step in HSV entry as well.

**Figure 3.**
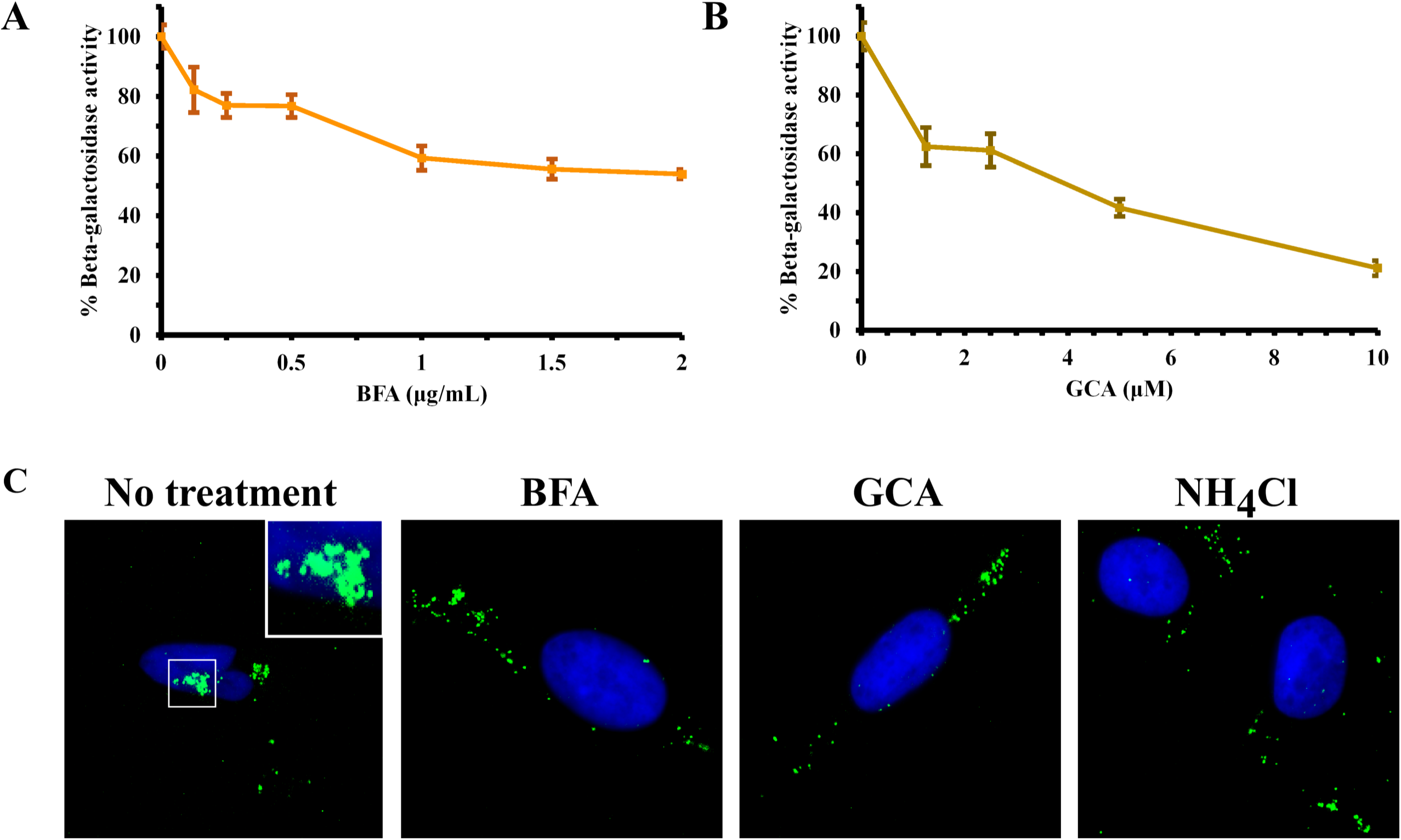
Golgi-inhibitors brefeldin A and Golgicide A impair HSV entry. CHO-HVEM cells were pretreated for 20 min with the indicated concentrations of (A) BFA or (B) GCA. HSV-1 strain KOS was added (MOI of 1) for 6 h in the constant presence of drug. Beta-galactosidase activity at 595 nm is indicative of successful HSV entry. Beta-galactosidase activity in the absence of drug was set to 100%. (C) CHO-HVEM cells on coverslips were infected with HSV-1 K26GFP (MOI of 100) and concurrently exposed to 2 µg/ml BFA, 10 µM GCA or 50 mM NH4Cl for 2.5 h. Cells were fixed, and nuclei were stained with DAPI (blue). In untreated cells, the majority of incoming capsids (green) were detected at or near the nucleus. In BFA, GCA, and control NH4Cl, treated cells, capsids were trapped in the cell periphery.

### Role of Rab9 and Rab11 in HSV-1 entry

Since retrograde transport and/or the TGN play a potential role in endocytic entry of HSV, we investigated Rab GTPases that control retrograde transport to the TGN. Rab9 mediates retrograde transport from the late endosome to the TGN (46, 47), and Rab11 controls retrograde transport from early/recycling endosomes-to-TGN (EE/RE-to-TGN) (50, 72). Other viruses take advantage of the recycling (45, 77) and TGN (57, 78) compartments. To test the roles of Rab9 or Rab11 in HSV entry, CHO-HVEM cells were treated with appropriate siRNAs. Knockdown of Rab9 or Rab11 was confirmed by Western blot (Fig. 4A). As determined by densitometry, Rab9 and Rab11 were reduced by 75 and 83%, respectively. As further confirmation, in Rab9 or Rab11-depleted cells, fluorescently labelled LysoTracker or transferrin, respectively was detected in enlarged vesicles relative to control cells (Fig. 2A-B). HSV-1 KOS was added to cells (MOI of 3) and infectivity was measured by indirect immunofluorescence at 6 hr p.i. (Fig. 4B). Silencing of Rab9 or Rab11 resulted in a slight decrease in HSV-1 infectivity (Fig. 4B). Under the conditions tested, the results suggest that these host cell proteins may play a small role in HSV entry. There is also likely significant entry that occurs independent of Rab9 and Rab11 functions. A TGN role in HSV entry that is independent of these Rab GTPases is also possible.

**Figure 4.**
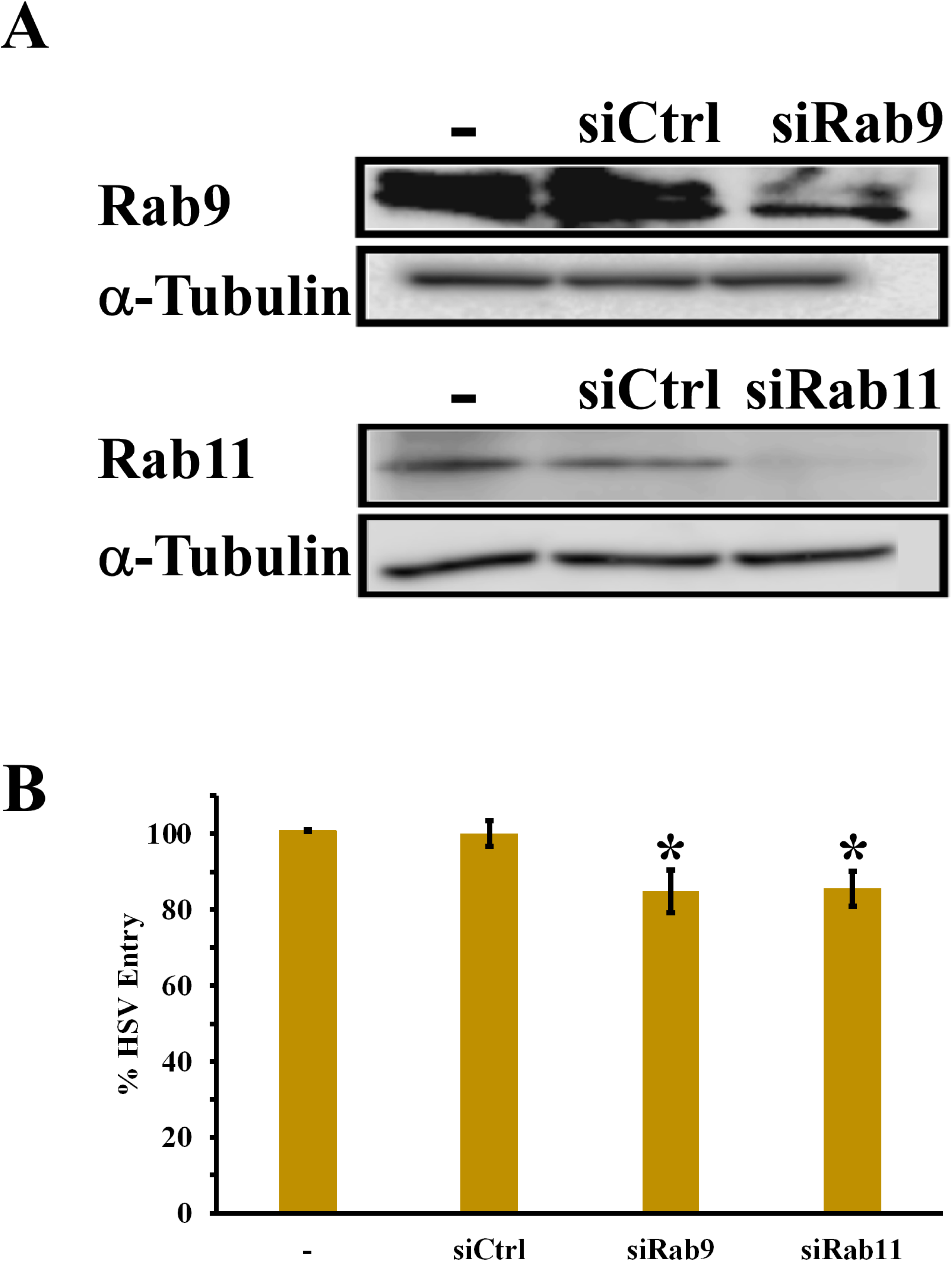
Effects of Rab9 and Rab11 knockdown on HSV entry and infection. (A) The knockdown efficiency of Rab9 or Rab11 was determined by Western blotting. CHO-HVEM cells (-) were transiently transfected with either control, Rab9 or Rab11 siRNAs for 24 h. Cell lysates were reacted with anti-Rab9 or anti-Rab11 antibody specific for the indicated proteins. α-Tubulin was used as an internal loading control. (B) CHO-HVEM cells (-) were transiently transfected with control, Rab9 or Rab11 siRNAs for 24 h. Control, Rab9, or Rab11 siRNA-treated cells were infected with HSV (MOI of 3). At 6 h p.i, infection was detected by quantitating HSV ICP4-positive cells via indirect immunofluorescence microscopy. At least 500 cells per cover slip were counted. Values are the mean ± SE of three independent experiments *P≤0.05.

A small molecule inhibitor was tested to probe further the role of retrograde trafficking in HSV entry. Retro-2 blocks retrograde traffic from EE to the TGN, and consequently inhibits cytosolic uptake of Shiga and ricin toxins (58, 79-82). Micromolar concentrations of retro-2 inhibits entry of human papillomavirus 16, BK virus, JC virus, and SV40, and transduction by adeno-associated virus serotype 2, all of which rely on retrograde transport mechanisms (58, 73, 75). Retro-2 treatment of CHO-HVEM cells had no inhibitory effect on HSV-1 entry as measured by the beta-galactosidase reporter assay (Fig. 5A). As a control for retro-2 activity, the effect on plaque formation was assessed. Retro 2.1, a derivative of retro-2, inhibits HSV-2 plaque formation on Vero cells, likely at the stage of assembly or egress (83). Retro-2 was effective at inhibiting HSV-1 infectivity as measured by plaque formation (Fig. 5B), suggesting that retro-2 can exhibit inhibitory activity in our hands. Retro-2 inhibits the wrapping and egress of vaccinia virus particles, suggesting that the retrograde pathway is important for these steps of the vaccinia replication cycle (84). HSV entry is insensitive to retro-2 yet sensitive to inhibition by both BFA and GCA. This suggests that HSV entry is distinct from Shiga toxin trafficking, although it may share overlapping features.

**Figure 5.**
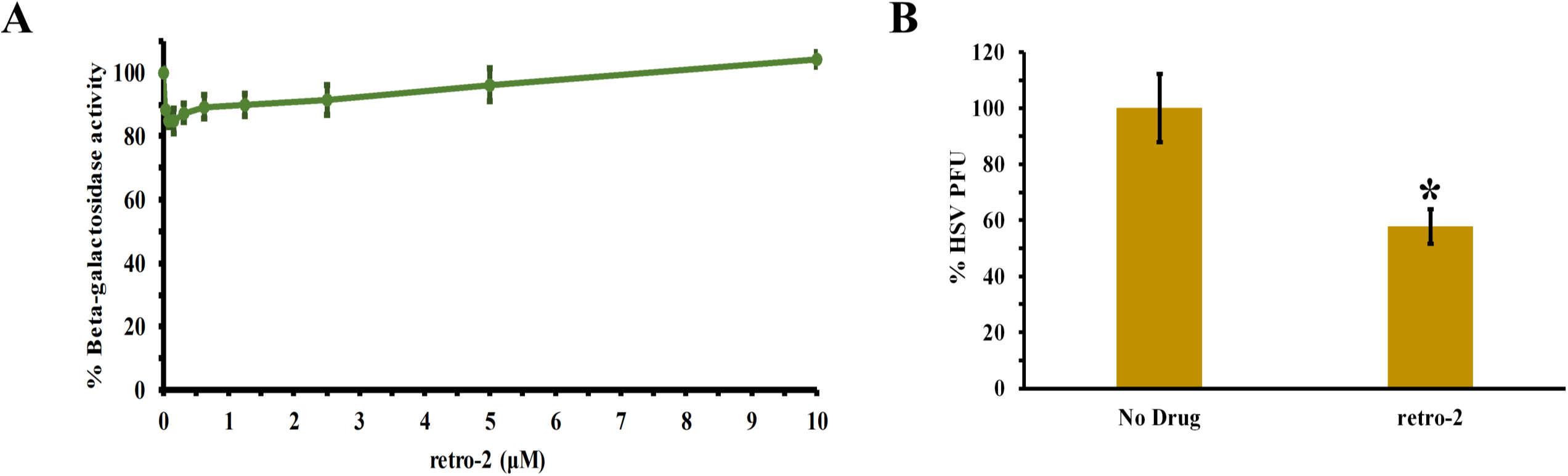
Effects of Retro-2 on HSV entry and infection. (A) CHO-HVEM cells were pretreated with the indicated concentrations of retro-2 for 4 h. HSV-1 strain KOS was added (MOI ∼ 1) for 6 h in the presence of drug. Beta-galactosidase activity in the absence of drug was set to 100% entry. (B) Vero cells were pretreated for 4 h with 50 μM retro-2. HSV-1 KOS (100 PFU/well) was added for 24 h at 37°C. Plaque formation in the absence of drug was set to 100%. Each experiment was performed in quadruplicate. Data shown are the averages ± SE of three independent experiments *P≤0.05.

The retrograde transport system is complex and selective. It relies on numerous tethering factors, small GTPases, and SNARES (85, 86). Other Rab proteins like Rab 6a, Rab 6IP, Rab 30 and the Rab 7b isoform mediate retrograde traffic from the endocytic compartment to the TGN (59, 87). The overlapping and varied functions of distinct isoforms of the same Rab protein (88) reflects the complexity of processes in which these proteins are involved (28, 89, 90). Notably, retrograde trafficking occurs from different endocytic compartments (EE, RE, LE) to the TGN and to the Golgi apparatus. There is a rapidly expanding network of proteins and processes involved in this interplay (91). The retromer, a conserved cytoplasmic protein complex, plays a central role in retrograde transport from endosome-to-Golgi, as well as from endosome-to-plasma membrane (57, 92, 93). HPV16 directly exploits the retromer at the early or late endosome and traffics to the TGN/Golgi via the retrograde pathway during cell entry (58). In contrast to HSV-1 (Fig. 5A), HPV16 entry is blocked by retro-2 in a concentration-dependent manner.

Overall this study suggests that HSV-1 entry by endocytosis is independent of the Rab GTPases that govern the conventional EE to LE to lysosome pathway. TGN function might be involved during HSV-1 low pH endocytosis entry in epithelial cells. It will be important to confirm the present results in more physiologically relevant cell types such as primary human keratinocytes. GCA, which blocks retrograde trafficking to the TGN, reduced HSV-1 infection of CHO-receptor cells by ∼ 79%. Further research is necessary to define the GCA- and BFA-sensitive nature of HSV entry. Importantly, alternate endocytic entry pathways may also function, even in the same cell type. The results reported here provide context for further characterization of HSV-1 entry by endocytosis and set the stage for identification of the intracellular site of membrane fusion.

## MATERIALS AND METHODS

### Cells

CHO-HVEM (M1A) cells (94), provided by R. Eisenberg and G. Cohen, University of Pennsylvania, are stably transformed with the human HVEM (or HveA) gene and contain the Escherichia coli *lacZ* gene under the control of the HSV-1 ICP4 gene promoter. CHO-HVEM cells were propagated in Ham’s F12 nutrient mixture (Gibco/Life Technologies) supplemented with 10% FBS, 150 μg of puromycin (Sigma-Aldrich, St. Louis, MO, USA)/ml, and 250 μg of G418 sulfate (Thermo Fisher Scientific, Fair Lawn, NJ, USA)/ml. Cells were sub-cultured in nonselective medium prior to use in all experiments. Vero cells (American Type Culture Collection, Manassas, VA) were propagated in Dulbecco’s modified Eagle’s medium (DMEM) (ThermoFisher Scientific, Grand Island, NY) supplemented with 10% fetal bovine serum (Atlanta Biologicals, Atlanta, GA), respectively. Baby hamster kidney (BHK) cells were propagated in DMEM high nutrient mixture (Gibco/Life Technologies) supplemented with 5% fetal bovine serum.

### Viruses

HSV-1 strain KOS (95) (provided by Priscilla Schaffer, Harvard Medical School) was propagated and titered on Vero cells. Vesicular stomatitis virus (VSV) Indiana strain, provided by Douglas Lyles, Wake Forest University, was titered on BHK cells and CHO-HVEM cells. HSV-1 KOS K26 GFP contains green fluorescent protein (GFP) fused to the N-terminus of the VP26 capsid protein (96) (provided by Prashant Desai, Johns Hopkins University). It was propagated and titrated on Vero cells.

### Transient transfection

CHO-HVEM cells were grown in 24-well plates to 70% confluence. Culture medium was removed, and the cells were transfected with siCtrl (sc-37007), siRab5A (sc-36344), siRab7A (sc-29460), siRab9A (sc-41831) or siRab11A (sc-36341) (Santa Cruz Biotechnology) using Lipo3000 transfection reagent (Thermo Fisher). Briefly, 60 pmol of siRNA was mixed with 1.5 μl Lipo3000 in 50 μl of serum-free OPTIMEM (Gibco Life Technologies) for 15 min at room temperature. Serum-free OPTIMEM medium was added to a final volume of 250 μl, and the transfection reaction was added to cells for 5 h at 37°C. The transfection mixture was replaced with medium supplemented with 10% FBS, and cultures were incubated for 24 or 48 h at 37°C.

### SDS-PAGE and Western blotting

At 24 or 48 h post-transfection, protein cell extracts were prepared by RIPA lysis buffer (50 mM Tris-HCl, EDTA 2 mM,150 mM NaCl, and 1% NP-40; pH 8) supplemented with cOmplete, Mini, EDTA-free Protease Inhibitor Cocktail (Roche). Total protein concentration was calculated with the BCA Protein Assay Kit (Pierce). Cell lysates were separated by SDS-PAGE on 4-20% Tris-glycine gels (Novex). Following transfer to nitrocellulose by electroblotting, membranes were blocked and incubated with antibodies to α-Tubulin (Sigma T9026), Rab5 (Abcam ab18211), Rab7 (Abcam ab137029), Rab9 (Invitrogen MA3-067), or Rab11 (Invitrogen 71-5300). Secondary antibody conjugated with horseradish peroxidase (Sigma) was added, followed by enhanced chemiluminescence substrate (SuperSignal West Dura Extended Duration, Thermo Scientific).

### HSV infectivity measured by immunofluorescence microscopy

At 24 or 48 h post-transfection, HSV-1 KOS (MOI of 3) or VSV (MOI of 0.5) was added to cell monolayers grown on glass coverslips in 24-well culture dishes. At 6 h p.i., cultures were washed with PBS, fixed in ice-cold methanol and blocked with 1% BSA in PBS. Anti-ICP4 MAb H1A021 or monoclonal Ab to VSV glycoprotein G (P5D4, Sigma) was added for 1 h followed by Alexa Fluor 488-labeled goat anti-mouse IgG (Invitrogen) for 30 min. Nuclei were counterstained with 5 ng/ml of DAPI. Coverslips were mounted on slides with Fluoromount G and visualized with a Leica DMi8 Fluorescence microscope at 10X magnification. Cells were counted with ImageJ. Three independent experiments were performed with three replicates per experiment.

### Cells transfected with dominant-negative Rab GTPase plasmids

Dominant-negative Rab5 (S34N) or Rab7 (T22N) or mock plasmids expressing the sequence of interest and Turbo RFP (Red Fluorescent Protein) gene under control of elongation factor, and the hygromycin-resistance gene under control of a mouse phosphoglycerate kinase 1 promoter were synthesized and sequenced by VectorBuilder (Chicago, Illinois). CHO-HVEM cells in a 25 cm^2^ flask were grown to 70% confluence and transfected with 7.5 μg of DNA using the Lipofectamine 3000 Transfection Kit (Invitrogen) according to the manufacturer’s instructions. At 5 h post-transfection, the transfection mixture was replaced with complete F12 medium. At 24 h post-transfection, medium was replaced with complete F12 supplemented with 500 μg/ml Hygromycin B (Invitrogen). When cells in a control untransfected flask were killed by Hygromycin B (∼72 h), the transfected cells were subcultured in selection medium. The stably transfected cells were passaged > 6 times under constant selection prior to use in viral entry experiments. For experiments, transfected or mock-transfected CHO-HVEM cells in Hygromycin B were plated and kept under Hygromycin selection until attachment. Following cell attachment, selective medium was removed and replaced with complete medium for 24 h.

### VSV plaque assay

Transfected cells in Hygromycin B were added to 6-well plates. Following cell attachment, selective medium was removed and replaced with complete medium. After 24 h, VSV was added and titered by limiting dilution. At 1 h p.i., the inoculum was removed and replaced with warm 2 ml 2% carboxymethyl cellulose in F12 medium with 2% FBS (CMC). At 48 h p.i., 2 ml of 10% formalin was layered onto the CMC for 2 h at room temperature. The overlay was removed, and fixed cells were washed with water. Crystal violet was added for 30 min at room temperature. Cells were rinsed with water. Plaques were visualized with a Leica stereoscope.

### Beta-galactosidase reporter assay of HSV-1 entry

HSV-1 KOS was added (MOI ∼ 1) in the continued presence of drugs. At 6 hr p.i., 0.5% IGEPAL (Sigma–Aldrich) was added to lyse the cells. Chlorophenol red-beta-D-galactopyranoside (Roche Diagnostics, Indianapolis, IN) substrate was added to cell lysates, and the beta-galactosidase activity was read at 595 nm with an ELx808 microtiter plate reader (BioTek Instruments, Winooski, VT, United States). Beta-galactosidase activity indicates successful viral entry (97).

### Transferrin and LysoTracker uptake assays

CHO-HVEM cells were grown in 24-well plates to 70% confluence and then transfected with siRab5A, siRab7A, siRab9A, or siRab11A as described above. For transferrin uptake, at 24 h post-transfection, cultures were washed twice and replenished with serum-free F12 medium for 80 min total at 37°C replacing the medium every 40 min. Cultures were incubated on ice at 4°C for 40 min. Transferrin Texas Red Conjugate (100 μg/ml) (Invitrogen T2875) was added to the wells for 50 min at 4°C. Cells were washed to remove unbound transferrin, and then serum-containing F12 medium was added at 37°C for 10 min. For LysoTracker uptake, at 24 h post-transfection, 100 nM LysoTracker Red DND-99 (Invitrogen L7528) in complete F12 medium was added for 30 min at 37°C. Cultures were washed twice with PBS and fixed with 3% paraformaldehyde–PBS at 37°C for 10 min. Fixed cells were washed twice with PBS, quenched with 50 mM NH4Cl at room temperature for 10 min, and then permeabilized with 0.1% Triton X-100 in PBS for 10 min. Nuclei were counterstained with 5 ng/ml of 4,6–diamidino-2-phenylindole dihydrochloride (DAPI; Roche). Coverslips were mounted on slides with Fluoromount G (Electron Microscopy Sciences, Hatfield, PA) and visualized with a Leica DMi8 fluorescence microscope at 100X magnification.

### Effect of Golgi inhibitors on HSV-1 entry

Confluent cell monolayers grown in 96-well plates were pretreated with a range of concentrations of brefeldin A (BFA, EMD Millipore CAS 20350-15-6) or Golgicide A (GCA, EMD Millipore CAS 1139889-93-2) for 20 min or with retro-2 (Sigma CAS 1429192-00-6) for 4 h at 37°C. Control samples were treated with vehicle DMSO. HSV-1 KOS was added and method for beta-galactosidase reporter assay was followed.

### Subcellular localization of entering GFP-tagged HSV

Sub-confluent CHO-HVEM cell monolayers grown on coverslips in 24-well plates were treated with 2 μg/ml BFA, 10 μM GCA or 50 mM NH_4_Cl in the presence of 0.5 mM cycloheximide for 20 min at 37°C. HSV-1 K26GFP (MOI of 100) was added for 2.5 h in the continued presence of agents. Cultures were washed twice with PBS and fixed with 3% paraformaldehyde–PBS at 37°C for 10 min. Fixed cells were washed twice with PBS, quenched with 50 mM NH4Cl at room temperature for 10 min, and then permeabilized with 0.1% Triton X-100 in PBS for 10 min. Nuclei were counterstained with 5 ng/ml of 4,6–diamidino-2-phenylindole dihydrochloride (DAPI; Roche). Coverslips were mounted on slides with Fluoromount G (Electron Microscopy Sciences, Hatfield, PA) and visualized with a Leica DMi8 fluorescence microscope at 100X magnification.

### HSV plaque assay

CHO-HVEM cells in 24 well plates were pre-treated with 50 μM Retro-2 (Sigma) for 4 h at 37°C. HSV-1 KOS (∼100 PFU/well) was added in the continued presence of drug. At 24 h p.i., cells were fixed with ice-cold methanol and acetone (2:1 ratio) for 20 min at −20°C and air-dried and assayed for HSV plaque formation. Cells were stained with rabbit polyclonal antibody to HSV, HR50 (Fitzgerald Industries, Concord, MA), washed thrice with PBS and incubated with 1:200 dilution of goat anti-rabbit IgG conjugated with horseradish peroxidase (Thermo Fisher Scientific) for 1 h at room temperature. Following three washes with PBS, 4-chloro-1-naphtol (Sigma) substrate was added and plaques were visualized with a Leica stereoscope.

### Statistical analysis

Student’s t-test with one tail distribution was used for infectivity experiments. Data are shown as geometric means with errors bars representing ± s.e.m. Significance is indicated as *P<0.05,**P<0.00005.

## ACKNOWLEDGMENTS

This work was supported by Public Health Service grant AI119159 (A.V.N.) from the National Institute of Allergy and Infectious Diseases. We thank Gary Cohen, Roselyn Eisenberg, Douglas Lyles, and Priscilla Schaffer for generous gifts of reagents. We thank Massaro Ueti for the use of the fluorescence microscope and members of the Nicola laboratory for helpful discussions and suggestions.

## REFERENCES

1. Roizman B, Knipe, DM., Whitley, RJ. 2013. Herpes simplex viruses, 6th ed. Lippincott Williams & Wilkins,.

2. Kennedy PG, Steiner I. 2013. Recent issues in herpes simplex encephalitis. J Neurovirol 19:346–350.

3. Liesegang TJ. 2001. Herpes simplex virus epidemiology and ocular importance. Cornea 20:1–13.

4. Nicola AV, McEvoy AM, Straus SE. 2003. Roles for endocytosis and low pH in herpes simplex virus entry into HeLa and Chinese hamster ovary cells. Journal of virology 77:5324–5332.

5. Nicola AV, Hou J, Major EO, Straus SE. 2005. Herpes simplex virus type 1 enters human epidermal keratinocytes, but not neurons, via a pH-dependent endocytic pathway. J Virol 79:7609–7616.

6. Sayers CL, Elliott G. 2016. Herpes Simplex Virus 1 Enters Human Keratinocytes by a Nectin-1-Dependent, Rapid Plasma Membrane Fusion Pathway That Functions at Low Temperature. J Virol 90:10379–10389.

7. Lycke E, Hamark B, Johansson M, Krotochwil A, Lycke J, Svennerholm B. 1988. Herpes simplex virus infection of the human sensory neuron. An electron microscopy study. Arch Virol 101:87–104.

8. Koyama AH, Uchida T. 1989. The effect of ammonium chloride on the multiplication of herpes simplex virus type 1 in Vero cells. Virus Res 13:271–281.

9. Nicola AV. 2016. Herpesvirus Entry into Host Cells Mediated by Endosomal Low pH. Traffic 17:965–975.

10. Nicola AV, Straus SE. 2004. Cellular and viral requirements for rapid endocytic entry of herpes simplex virus. Journal of virology 78:7508–7517.

11. Pastenkos G, Lee B, Pritchard SM, Nicola AV. 2018. Bovine Herpesvirus 1 Entry by a Low-pH Endosomal Pathway. J Virol 92.

12. Miller JL, Weed DJ, Lee BH, Pritchard SM, Nicola AV. 2019. Low-pH Endocytic Entry of the Porcine Alphaherpesvirus Pseudorabies Virus. J Virol 93.

13. Dollery SJ, Delboy MG, Nicola AV. 2010. Low pH-induced conformational change in herpes simplex virus glycoprotein B. J Virol 84:3759–3766.

14. Weed DJ, Nicola AV. 2017. Herpes simplex virus Membrane Fusion. Adv Anat Embryol Cell Biol 223:29–47.

15. Devadas D, Koithan T, Diestel R, Prank U, Sodeik B, Dohner K. 2014. Herpes simplex virus internalization into epithelial cells requires Na+/H+ exchangers and p21-activated kinases but neither clathrin- nor caveolin-mediated endocytosis. J Virol 88:13378–13395.

16. Clement C, Tiwari V, Scanlan PM, Valyi-Nagy T, Yue BYJT, Shukla D. 2006. A novel role for phagocytosis-like uptake in herpes simplex virus entry. The Journal of cell biology 174:1009–1021.

17. Petermann P, Haase I, Knebel-Morsdorf D. 2009. Impact of Rac1 and Cdc42 signaling during early herpes simplex virus type 1 infection of keratinocytes. J Virol 83:9759–9772.

18. Rahn E, Petermann P, Hsu MJ, Rixon FJ, Knebel-Morsdorf D. 2011. Entry pathways of herpes simplex virus type 1 into human keratinocytes are dynamin- and cholesterol-dependent. PloS one 6:e25464.

19. Nicola AV, Aguilar HC, Mercer J, Ryckman B, Wiethoff CM. 2013. Virus entry by endocytosis. Adv Virol 2013:469538.

20. Fuchs R, Ellinger, A., Pavelka, M., Peterlik, M., and Mellman, I. 1987. Endocytic coated vesicles do not exhibit ATP-dependent acidification in vitro. J Cell Biol 105.

21. Klumperman J, Raposo G. 2014. The complex ultrastructure of the endolysosomal system. Cold Spring Harbor perspectives in biology 6:a016857–a016857.

22. van Weering JRT, Verkade P, Cullen PJ. 2012. SNX-BAR-Mediated Endosome Tubulation is Co-ordinated with Endosome Maturation. Traffic 13:94–107.

23. Cossart P, Helenius A. 2014. Endocytosis of viruses and bacteria. Cold Spring Harb Perspect Biol 6.

24. Yamauchi Y, Helenius A. 2013. Virus entry at a glance. J Cell Sci 126:1289–1295.

25. Staring J, Raaben M, Brummelkamp TR. 2018. Viral escape from endosomes and host detection at a glance. J Cell Sci 131.

26. Lozach PY, Huotari J, Helenius A. 2011. Late-penetrating viruses. Curr Opin Virol 1:35–43.

27. Campadelli-Fiume G, Arsenakis M, Farabegoli F, Roizman B. 1988. Entry of herpes simplex virus 1 in BJ cells that constitutively express viral glycoprotein D is by endocytosis and results in degradation of the virus. J Virol 62:159–167.

28. Pylypenko O, Hammich H, Yu IM, Houdusse A. 2018. Rab GTPases and their interacting protein partners: Structural insights into Rab functional diversity. Small GTPases 9:22–48.

29. Ao X, Zou L, Wu Y. 2014. Regulation of autophagy by the Rab GTPase network. Cell Death Differ 21:348–358.

30. Hutagalung AH, Novick PJ. 2011. Role of Rab GTPases in membrane traffic and cell physiology. Physiol Rev 91:119–149.

31. Stenmark H, Olkkonen VM. 2001. The Rab GTPase family. Genome Biol 2:REVIEWS3007.

32. Elgner F, Hildt E, Bender D. 2018. Relevance of Rab Proteins for the Life Cycle of Hepatitis C Virus. Front Cell Dev Biol 6:166.

33. De Franceschi N, Hamidi H, Alanko J, Sahgal P, Ivaska J. 2015. Integrin traffic -the update. J Cell Sci 128:839–852.

34. Militello R, Colombo MI. 2013. Small GTPases as regulators of cell division. Commun Integr Biol 6:e25460.

35. Numrich J, Ungermann C. 2014. Endocytic Rabs in membrane trafficking and signaling. Biol Chem 395:327–333.

36. van Ijzendoorn SC, Mostov KE, Hoekstra D. 2003. Role of rab proteins in epithelial membrane traffic. Int Rev Cytol 232:59–88.

37. Spearman P. 2018. Viral interactions with host cell Rab GTPases. Small GTPases 9:192–201.

38. Gorvel JP, Chavrier P, Zerial M, Gruenberg J. 1991. rab5 controls early endosome fusion in vitro. Cell 64:915–925.

39. Bucci C, Parton RG, Mather IH, Stunnenberg H, Simons K, Hoflack B, Zerial M. 1992. The small GTPase rab5 functions as a regulatory factor in the early endocytic pathway. Cell 70:715–728.

40. McLauchlan H, Newell J, Morrice N, Osborne A, West M, Smythe E. 1998. A novel role for Rab5-GDI in ligand sequestration into clathrin-coated pits. Curr Biol 8:34–45.

41. Feng Y, Press B, Wandinger-Ness A. 1995. Rab 7: an important regulator of late endocytic membrane traffic. J Cell Biol 131:1435–1452.

42. Vitelli R, Santillo M, Lattero D, Chiariello M, Bifulco M, Bruni CB, Bucci C. 1997. Role of the small GTPase Rab7 in the late endocytic pathway. J Biol Chem 272:4391–4397.

43. Gutierrez MG, Munafo DB, Beron W, Colombo MI. 2004. Rab7 is required for the normal progression of the autophagic pathway in mammalian cells. J Cell Sci 117:2687–2697.

44. Jager S, Bucci C, Tanida I, Ueno T, Kominami E, Saftig P, Eskelinen EL. 2004. Role for Rab7 in maturation of late autophagic vacuoles. J Cell Sci 117:4837–4848.

45. Liu C-C, Zhang Y-N, Li Z-Y, Hou J-X, Zhou J, Kan L, Zhou B, Chen P-Y. 2017. Rab5 and Rab11 Are Required for Clathrin-Dependent Endocytosis of Japanese Encephalitis Virus in BHK-21 Cells. Journal of virology 91:e01113–01117.

46. Lombardi D, Soldati T, Riederer MA, Goda Y, Zerial M, Pfeffer SR. 1993. Rab9 functions in transport between late endosomes and the trans Golgi network. EMBO J 12:677–682.

47. Dintzis SM, Pfeffer SR. 1990. The mannose 6-phosphate receptor cytoplasmic domain is not sufficient to alter the cellular distribution of a chimeric EGF receptor. EMBO J 9:77–84.

48. Zhang Y-N, Liu Y-Y, Xiao F-C, Liu C-C, Liang X-D, Chen J, Zhou J, Baloch AS, Kan L, Zhou B, Qiu H-J. 2018. Rab5, Rab7, and Rab11 Are Required for Caveola-Dependent Endocytosis of Classical Swine Fever Virus in Porcine Alveolar Macrophages. Journal of virology 92:e00797–00718.

49. Ullrich O, Reinsch S, Urbe S, Zerial M, Parton RG. 1996. Rab11 regulates recycling through the pericentriolar recycling endosome. J Cell Biol 135:913–924.

50. Wilcke M, Johannes L, Galli T, Mayau V, Goud B, Salamero J. 2000. Rab11 regulates the compartmentalization of early endosomes required for efficient transport from early endosomes to the trans-golgi network. J Cell Biol 151:1207–1220.

51. Welz T, Wellbourne-Wood J, Kerkhoff E. 2014. Orchestration of cell surface proteins by Rab11. Trends Cell Biol 24:407–415.

52. Gianni T, Campadelli-Fiume G, Menotti L. 2004. Entry of herpes simplex virus mediated by chimeric forms of nectin1 retargeted to endosomes or to lipid rafts occurs through acidic endosomes. J Virol 78:12268–12276.

53. Campadelli-Fiume G, Cocchi F, Menotti L, Lopez M. 2000. The novel receptors that mediate the entry of herpes simplex viruses and animal alphaherpesviruses into cells. Rev Med Virol 10:305–319.

54. Sieczkarski SB, Whittaker GR. 2003. Differential requirements of Rab5 and Rab7 for endocytosis of influenza and other enveloped viruses. Traffic 4:333–343.

55. Quirin K, Eschli B, Scheu I, Poort L, Kartenbeck J, Helenius A. 2008. Lymphocytic choriomeningitis virus uses a novel endocytic pathway for infectious entry via late endosomes. Virology 378:21–33.

56. Liebl D, Difato F, Hornikova L, Mannova P, Stokrova J, Forstova J. 2006. Mouse polyomavirus enters early endosomes, requires their acidic pH for productive infection, and meets transferrin cargo in Rab11-positive endosomes. J Virol 80:4610–4622.

57. DiMaio D, Burd CG, Goodner K. 2015. Riding the R Train into the Cell. PLoS Pathog 11:e1005036.

58. Lipovsky A, Popa A, Pimienta G, Wyler M, Bhan A, Kuruvilla L, Guie MA, Poffenberger AC, Nelson CD, Atwood WJ, DiMaio D. 2013. Genome-wide siRNA screen identifies the retromer as a cellular entry factor for human papillomavirus. Proc Natl Acad Sci U S A 110:7452–7457.

59. Sapp MJ. 2013. HPV virions hitchhike a ride on retromer complexes. Proc Natl Acad Sci U S A 110:7116–7117.

60. Casey JR, Grinstein S, Orlowski J. 2010. Sensors and regulators of intracellular pH. Nat Rev Mol Cell Biol 11:50–61.

61. Paroutis P, Touret N, Grinstein S. 2004. The pH of the secretory pathway: measurement, determinants, and regulation. Physiology (Bethesda) 19:207–215.

62. Demaurex N, Furuya W, D’Souza S, Bonifacino JS, Grinstein S. 1998. Mechanism of acidification of the trans-Golgi network (TGN). In situ measurements of pH using retrieval of TGN38 and furin from the cell surface. J Biol Chem 273:2044–2051.

63. Dollery SJ, Wright CC, Johnson DC, Nicola AV. 2011. Low-pH-dependent changes in the conformation and oligomeric state of the prefusion form of herpes simplex virus glycoprotein B are separable from fusion activity. J Virol 85:9964–9973.

64. Siekavizza-Robles CR, Dollery SJ, Nicola AV. 2010. Reversible conformational change in herpes simplex virus glycoprotein B with fusion-from-without activity is triggered by mildly acidic pH. Virol J 7:352.

65. Weed DJ, Pritchard SM, Gonzalez F, Aguilar HC, Nicola AV. 2017. Mildly Acidic pH Triggers an Irreversible Conformational Change in the Fusion Domain of Herpes Simplex Virus 1 Glycoprotein B and Inactivation of Viral Entry. J Virol 91.

66. Weed DJ, Dollery SJ, Komala Sari T, Nicola AV. 2018. Acidic pH Mediates Changes in Antigenic and Oligomeric Conformation of Herpes Simplex Virus gB and Is a Determinant of Cell-Specific Entry. J Virol 92.

67. Lippincott-Schwartz J, Yuan L, Tipper C, Amherdt M, Orci L, Klausner RD. 1991. Brefeldin A’s effects on endosomes, lysosomes, and the TGN suggest a general mechanism for regulating organelle structure and membrane traffic. Cell 67:601–616.

68. Huang S, Wang Y. 2017. Golgi structure formation, function, and post-translational modifications in mammalian cells. F1000Res 6:2050.

69. Saenz JB, Sun WJ, Chang JW, Li J, Bursulaya B, Gray NS, Haslam DB. 2009. Golgicide A reveals essential roles for GBF1 in Golgi assembly and function. Nat Chem Biol 5:157–165.

70. Klausner RD, Donaldson JG, Lippincott-Schwartz J. 1992. Brefeldin A: insights into the control of membrane traffic and organelle structure. J Cell Biol 116:1071–1080.

71. Lippincott-Schwartz J, Yuan LC, Bonifacino JS, Klausner RD. 1989. Rapid redistribution of Golgi proteins into the ER in cells treated with brefeldin A: evidence for membrane cycling from Golgi to ER. Cell 56:801–813.

72. Mallard F, Tang BL, Galli T, Tenza D, Saint-Pol A, Yue X, Antony C, Hong W, Goud B, Johannes L. 2002. Early/recycling endosomes-to-TGN transport involves two SNARE complexes and a Rab6 isoform. J Cell Biol 156:653–664.

73. Nelson CD, Carney DW, Derdowski A, Lipovsky A, Gee GV, O’Hara B, Williard P, DiMaio D, Sello JK, Atwood WJ. 2013. A retrograde trafficking inhibitor of ricin and Shiga-like toxins inhibits infection of cells by human and monkey polyomaviruses. mBio 4:e00729–00713.

74. Bantel-Schaal U, Hub B, Kartenbeck J. 2002. Endocytosis of adeno-associated virus type 5 leads to accumulation of virus particles in the Golgi compartment. J Virol 76:2340–2349.

75. Nonnenmacher ME, Cintrat JC, Gillet D, Weber T. 2015. Syntaxin 5-dependent retrograde transport to the trans-Golgi network is required for adeno-associated virus transduction. J Virol 89:1673–1687.

76. Laniosz V, Dabydeen SA, Havens MA, Meneses PI. 2009. Human papillomavirus type 16 infection of human keratinocytes requires clathrin and caveolin-1 and is brefeldin a sensitive. J Virol 83:8221–8232.

77. Vale-Costa S, Amorim MJ. 2016. Recycling Endosomes and Viral Infection. Viruses 8:64.

78. Day PM, Thompson CD, Schowalter RM, Lowy DR, Schiller JT. 2013. Identification of a role for the trans-Golgi network in human papillomavirus 16 pseudovirus infection. J Virol 87:3862–3870.

79. Stechmann B, Bai SK, Gobbo E, Lopez R, Merer G, Pinchard S, Panigai L, Tenza D, Raposo G, Beaumelle B, Sauvaire D, Gillet D, Johannes L, Barbier J. 2010. Inhibition of retrograde transport protects mice from lethal ricin challenge. Cell 141:231–242.

80. Nicolas V, Lievin-Le Moal V. 2020. Small Trafficking Inhibitor Retro-2 Disrupts the Microtubule-Dependent Trafficking of Autophagic Vacuoles. Front Cell Dev Biol 8:464.

81. Forrester A, Rathjen SJ, Daniela Garcia-Castillo M, Bachert C, Couhert A, Tepshi L, Pichard S, Martinez J, Munier M, Sierocki R, Renard HF, Augusto Valades-Cruz C, Dingli F, Loew D, Lamaze C, Cintrat JC, Linstedt AD, Gillet D, Barbier J, Johannes L. 2020. Functional dissection of the retrograde Shiga toxin trafficking inhibitor Retro-2. Nat Chem Biol 16:327–336.

82. Farhan H. 2020. Rendezvous of Retro-2 at the ER. Nat Chem Biol 16:229–230.

83. Dai W, Wu Y, Bi J, Wang J, Wang S, Kong W, Barbier J, Cintrat JC, Gao F, Jiang Z, Gillet D, Su W, Jiang C. 2018. Antiviral Effect of Retro-2.1 against Herpes Simplex Virus Type 2 In Vitro. J Microbiol Biotechnol 28:849–859.

84. Sivan G, Weisberg AS, Americo JL, Moss B. 2016. Retrograde Transport from Early Endosomes to the trans-Golgi Network Enables Membrane Wrapping and Egress of Vaccinia Virus Virions. J Virol 90:8891–8905.

85. Bonifacino JS, Rojas R. 2006. Retrograde transport from endosomes to the trans-Golgi network. Nat Rev Mol Cell Biol 7:568–579.

86. Schindler C, Chen Y, Pu J, Guo X, Bonifacino JS. 2015. EARP is a multisubunit tethering complex involved in endocytic recycling. Nat Cell Biol 17:639–650.

87. Zulkefli KL. 2017. Role of Rab11 and Rab30 in regulating the endosome-trans-Golgi networkUniversity of Melbourne.

88. Schwartz SL, Cao C, Pylypenko O, Rak A, Wandinger-Ness A. 2007. Rab GTPases at a glance. J Cell Sci 120:3905–3910.

89. Pfeffer SR. 2005. Structural clues to Rab GTPase functional diversity. J Biol Chem 280:15485–15488.

90. Diekmann Y, Seixas E, Gouw M, Tavares-Cadete F, Seabra MC, Pereira-Leal JB. 2011. Thousands of rab GTPases for the cell biologist. PLoS Comput Biol 7:e1002217.

91. Johannes L, Popoff V. 2008. Tracing the retrograde route in protein trafficking. Cell 135:1175–1187.

92. Lu L, Hong W. 2014. From endosomes to the trans-Golgi network. Semin Cell Dev Biol 31:30–39.

93. Seaman MN. 2012. The retromer complex - endosomal protein recycling and beyond. J Cell Sci 125:4693–4702.

94. Montgomery RI, Warner MS, Lum BJ, Spear PG. 1996. Herpes simplex virus-1 entry into cells mediated by a novel member of the TNF/NGF receptor family. Cell 87:427–436.

95. Macdonald SJ, Mostafa HH, Morrison LA, Davido DJ. 2012. Genome sequence of herpes simplex virus 1 strain KOS. J Virol 86:6371–6372.

96. Desai P, Person S. 1998. Incorporation of the green fluorescent protein into the herpes simplex virus type 1 capsid. J Virol 72:7563–7568.

97. Delboy MG, Patterson JL, Hollander AM, Nicola AV. 2006. Nectin-2-mediated entry of a syncytial strain of herpes simplex virus via pH-independent fusion with the plasma membrane of Chinese hamster ovary cells. Virology Journal 3.

